# Typhoid toxin of *Salmonella* Typhi elicits host antimicrobial response during acute typhoid fever

**DOI:** 10.1101/2024.12.13.628371

**Authors:** Salma Srour, Mohamed ElGhazaly, Daniel O’Connor, Malick M Gibani, Thomas C Darton, Andrew J Pollard, Mark O Collins, Daniel Humphreys

## Abstract

*Salmonella* Typhi secretes typhoid toxin that activates cellular DNA damage responses (DDR) during acute typhoid fever. Human infection challenge studies revealed that the toxin suppresses bacteraemia via unknown mechanisms. Using quantitative proteomic analysis on the plasma of bacteraemic participants, we demonstrate that wild-type toxigenic *Salmonella* induced secretion of lysozyme (LYZ) and apolipoprotein C3 (APOC3). Recombinant typhoid toxin or infection with toxigenic *Salmonella* recapitulate LYZ and APOC3 secretion in cultured cells, which involved ATM/ATR-dependent DDRs and confirmed observations in typhoid fever. LYZ alone inhibited secretion of virulence effector proteins SipB and SopE. LYZ caused loss of *Salmonella* morphology characterised by spheroplast formation. Spheroplast formation was mediated by LYZ and enhanced by lactoferrin, which was identified by proteomics in participants with typhoid fever. Our findings may indicate that toxin-induced DDRs elicit antimicrobial responses, which suppress *Salmonella* bacteraemia during typhoid fever.

## Introduction

Acute typhoid fever is caused by *Salmonella enterica* serovar Typhi (11 million cases, 116,800 deaths per year), which is a major health problem disproportionately affecting low- and middle-income countries [1]. Typhoid fever is established when *S.*Typhi invades the intestinal mucosa from where the pathogen disseminates into the bloodstream resulting in an asymptomatic primary bacteraemia. Replication then occurs in lymphoid tissues and after an incubation period of 7-10 days a secondary bacteraemia coincides with febrile symptoms and the onset of acute enteric fever and shedding of transmissible bacteria in stool. A small proportion of individuals develop asymptomatic chronic *S.*Typhi carriage, further contributing to ongoing community transmission to new hosts [1]. The control of *S.*Typhi is possible through provision of clean water and vaccines, but hampered by inadequate diagnostics, and rising antimicrobial-resistance.

To initiate infections, *Salmonella* inject virulence effectors directly into host intestinal epithelial cells using a type 3 secretion system (T3SS) that mediates pathogen macropinocytosis [2]. This includes essential effectors such as SipB [3], which form a translocon in the host plasma membrane through which effectors such as SopE, SptP and SopB are translocated to manipulate Rho and Arf GTPase signalling [2, 4–7]. Following macropinocytosis, *S.*Typhi resides with a *Salmonella-*containing vacuole (SCV) from where it expresses the typhoid toxin comprising PltB-PltA-CdtB subunits that are exocytosed into the extracellular milieu [8]. Once deployed, the PltB subunit binds to sialylated glycans on host surface receptors facilitating toxin endocytosis [9]. Reduction of disulphide bonds linking PltA-CdtB liberates the toxigenic DNase1-like subunit CdtB, which translocates to the nucleus where it activates DDRs through nuclease activity [8–11].

The mechanisms by which toxin-mediated host DDRs influence host pathogen interactions in humans are unclear. The human-specificity of *S.*Typhi has meant reliance on infection of human cells *in vitro* or using non-typhoidal *Salmonella* in infections of mice. In human cells, toxin-induced DDRs causing DNA replication stress and senescence leading to release of a host secretome [8–10, 12, 13]. In mice, injection of purified toxin causes typhoid fever-like symptoms and fatality [9] while infection with non-typhoidal *Salmonella* (NTS) encoding typhoid toxin suppressed host damage and promoted chronic infections [14, 15]. To advance our understanding of typhoid toxin, typhoid fever was studied using a controlled human infection model, which involves deliberate infection of volunteers [1]. Human participants were challenged with either a wild-type (WT) *S.*Typhi strain expressing the toxin or a toxin-negative (TN) strain lacking the genes *pltB pltA* and *cdtB* [16]. Counterintuitively, disease tended to be more severe in participants infected with *S.*Typhi-TN (severe typhoid in 7% due to *S.*Typhi-WT; 27% with *S.*Typhi-TN), which was reflected by significantly prolonged bacteraemia relative to participants infected with *S.*Typhi-WT (WT, 48h; TN, 98h).

The findings in Gibani et al 2019 indicate that host responses to the typhoid toxin suppress the duration of bacteraemia but the mechanisms are not known[16]. Many studies relating to innate defences against *Salmonella* have focussed on murine non-typhoidal *Salmonella* (NTS) models due to their ability to infect mice. It is not clear how this translates to human-restricted *S.*Typhi infections, which, in contrast to NTS, limits activation of innate immune responses [17]. It is also not clear how host DDRs coordinate defences against pathogens themselves: attention has focussed on mechanisms by which pathogens disarm the host DDR for microbial benefit [18], how bacterial genotoxins contribute to cancer [19] or trigger immune signalling pathways [20]. It was thus hypothesised that host DDRs mount a defence against toxigenic pathogens such as *S.*Typhi, which could highlight how DDRs counteract bacterial pathogens. We sought to address whether toxin-mediated effects on the host proteome could explain the reduced duration of *S.*Typhi bacteraemia.

## Results

### Typhoid toxin manipulates the host secretome in bacteraemic humans with typhoid fever

Typhoid toxin induced a DDR-dependent host secretome in cultured fibroblasts, intestinal epithelial cells and macrophages [10, 12, 13]. Thus, it was hypothesised that in human participants challenged with WT toxigenic *S.*Typhi [16], toxin-induced secretion would be reflected in the host proteome. To identify proteomic signatures in response to typhoid toxin, we performed LC-MS/MS analysis on plasma harvested from bacteraemic participants at the time of typhoid diagnosis following infection with either WT (cyan) or TN (magenta) *S.*Typhi (**Fig 1A**). As a reference for toxin-induced effects, proteomics was also performed on the same participants prior to infection (baseline), i.e. 20 participants before infection and 20 participants after infection. Plasma consists of high abundance proteins (∼94%) conserved between individuals [21], which were first removed by immunodepletion to reduce the dynamic range of the plasma proteome. LC-MS/MS identified 641 proteins at a 1% FDR **(Dataset S1)**, and label-free quantification was used to measure differences in the abundance of proteins between groups of participants (**Dataset S1, S2**). After data filtering and normalisation, statistical analysis was performed on 440 proteins to identify significant differences between the groups using a permutation-based FDR of 0.05 **(Dataset S2)**.

**Figure 1:**
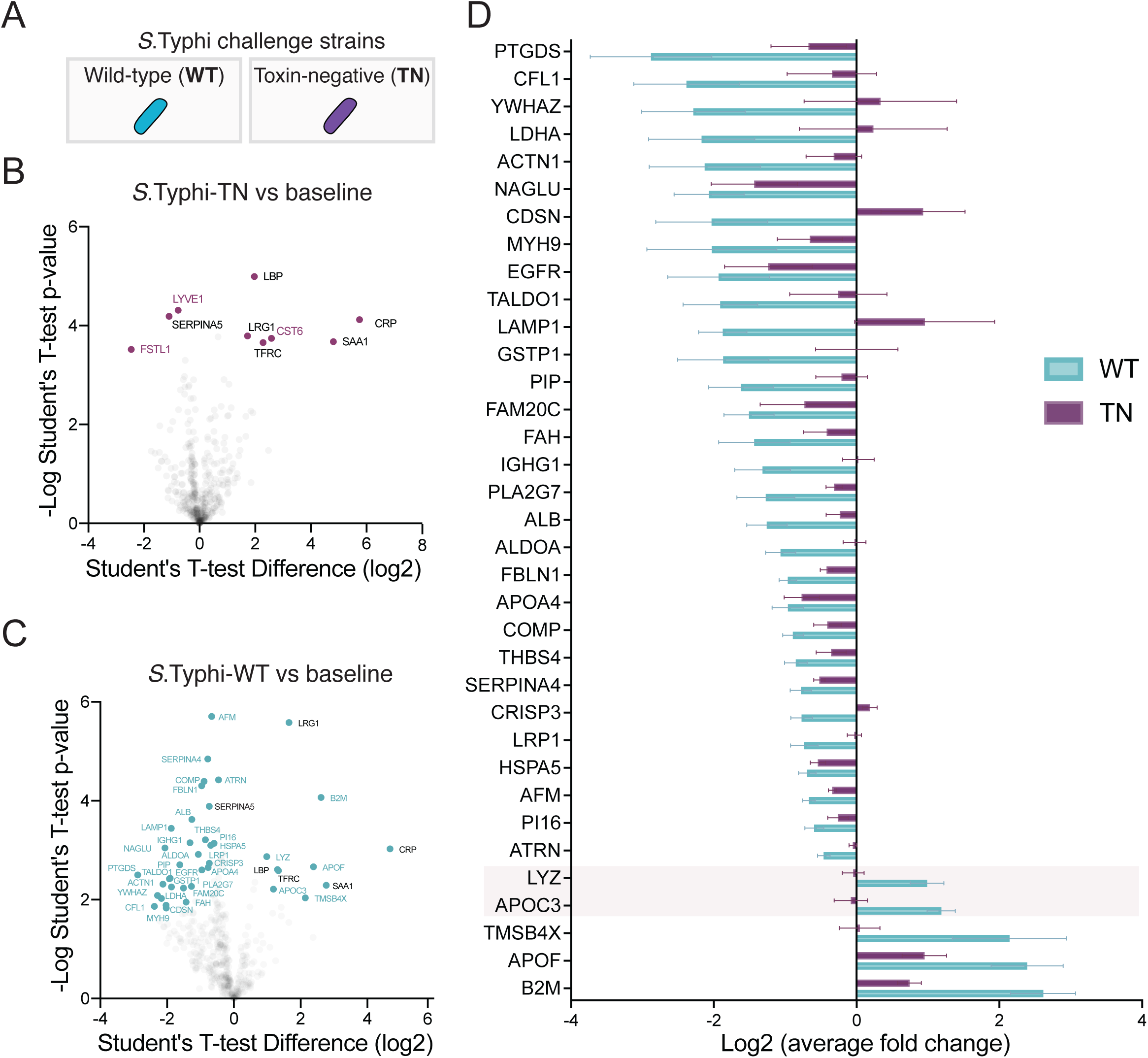
*Salmonella* Typhi exhibits a toxin induced secretome in host organisms. **(A)** Schematic showing the strains used in human infection challenge study, *S*. Typhi-WT (cyan) expressing typhoid toxin and toxin-negative (TN; *ΔpltA, ΔpltB, ΔcdtB*) *S*. Typhi (purple). Volcano plots from proteomics showing host responses in plasma to infection by **(B)** S. Typhi TN, or **(C)** S. Typhi WT in bacteraemic participants at TD (time of diagnosis) relative to uninfected baseline. Toxin-dependent proteins identified (cyan text), toxin-independent proteins (purple), and proteins identified in both analyses (black) are indicated. **(D)** Heatmap listing fold change of toxin-induced host proteins between WT (cyan) and TN (purple) *S.* Typhi-infected participants. LYZ and APOC3 highlighted.

In participants infected with TN *S.*Typhi, we found that 9 proteins were significantly different relative to baseline permutation-based FDR of 0.05 (**Fig 1B**). In contrast, 41 host proteins were differentially regulated in response to the typhoid toxin during acute typhoid fever (**Fig 1C**). Of the 9 TN-specific proteins, 6 proteins (CRP, LBP, LRG1, SAA1, TFRC) overlapped with the WT group identifying them as infection-specific and toxin-independent proteins (**Fig 1B**; black text). Only 3 proteins (CST6, FSTL1, LYVE1) were unique to the TN group marking them as TN-specific (**Fig 1B, 1D**; magenta). This contrasted with participants exposed to typhoid toxin as 35 proteins were enriched in the WT group (**Fig 1C, 1D**; cyan), consistent with a toxin-mediated effect on the host secretome. This included 5 WT-specific proteins of increased abundance: beta-2-microglobulin (B2M), apolipoprotein C3 (APOC3), apolipoprotein F (APOF), lysozyme (LYZ), and thymosin (TMSB4X) (**Fig 1D**), all of which are known secreted proteins [22]. The remaining 30 proteins in the WT group were of decreased abundance (**Fig 1D**). Taken together, the findings indicate that typhoid toxin manipulates the host proteome during acute typhoid fever.

### Typhoid toxin elicits secretion of APOC3 during acute typhoid fever

We next investigated whether changes in protein abundance could indicate why *S.*Typhi bacteraemia was prolonged in the absence of the toxin. To narrow our focus, we concentrated on the 5 WT-specific proteins of increased abundance in acute typhoid fever (**Fig 1D**: B2M, APOC3, APOF, LYZ, TMSB4X). B2M, APOF, and TMSB4X were most abundant but relative to *S.*Typhi-WT these proteins had also increased in response to *S.*Typhi-TN, albeit to a small extent with THSB4X. In contrast, APOC3 and LYZ increased in response to *S.*Typhi-WT in a toxin-dependent manner as both proteins were slightly reduced in response to *S.*Typhi-TN (**Fig 1D**: see highlight). LYZ is a ubiquitous 15kDa component of the innate immune response that hydrolyses β-1,4,glycosidic bonds in cell walls between *N*-acetylmuramic acid and *N*-acetylglucosamine in peptidoglycan causing bacterial lysis [23]. A role for Apolipoprotein C-III (APOC3) is less clear and required further investigation: APOC3 is a 9kDa apolipoprotein only expressed in the liver within hepatocytes and epithelial cells of the gastrointestinal tract, which increases the concentration of free lipids in the blood [24]. High concentrations of APOC3 are correlated with hypertriglyceridemia [24] but no role during bacterial infection is known.

When we studied APOC3 in CACO2 intestinal cells, we found that very little APOC3 was observed in untreated control cells (**Fig 2A, 2B**). In contrast, CACO2 cells treated for 2h with purified recombinant wild-type typhoid toxin (TxWT) expressed APOC3 at 96h (**Fig 2A, 2B**), which was observed in the nucleus as previously described [25]. APOC3 expression was coincident with activation of DDRs marked by γH2AX (**Fig 2A, 2C**), which corresponded to cell-cycle arrest as indicated by lack of DNA synthesis incorporating the nucleotide analogue EdU (**Fig 2A, 2D**). Indeed, APOC3 expression appeared dependent on DDRs as treating cells with typhoid toxin deficient in DNase activity due to its H160Q substitution (TxHQ) induced no γH2AX or APOC3 (**Fig 2A-C**), which was consistent with EdU-positive nuclei marking replicating cells (**Fig 2A, 2D**). Increased APOC3 expression in response to toxin-induced DDRs was mirrored by a relative increase in APOC3 secretion from the CACO2 cells at 96h (**Fig 2E**). In contrast, when we examined APOC3 inside intoxicated HepG2 liver cells, we found that γH2AX DDRs induced by TxWT or the drug aphidicolin, which inhibits DNA polymerase α, had no effect on APOC3 that was expressed equivalently in all conditions (**Fig S1A-C**). This indicated that increased APOC3 originated from infected intestinal epithelial cells during typhoid fever [16]. To test this possibility during infection, we examined APOC3 induction during infection with toxigenic *Salmonella* Javiana (**Fig 3**), a non-typhoidal *Salmonella* strain encoding typhoid toxin and was used for biosafety reasons. When CACO2 cells were infected with wild-type *S.*Javiana encoding typhoid toxin, APOC3 was observed in cells displaying increased levels of γH2AX relative to untreated and toxin-negative *ΔcdtB S.*Javiana (**Fig 3A-C**). In summary, we find that toxin-induced γH2AX-labelled DDRs modulates expression and secretion of APOC3, which was identified in human participants with acute typhoid fever.

**Figure 2:**
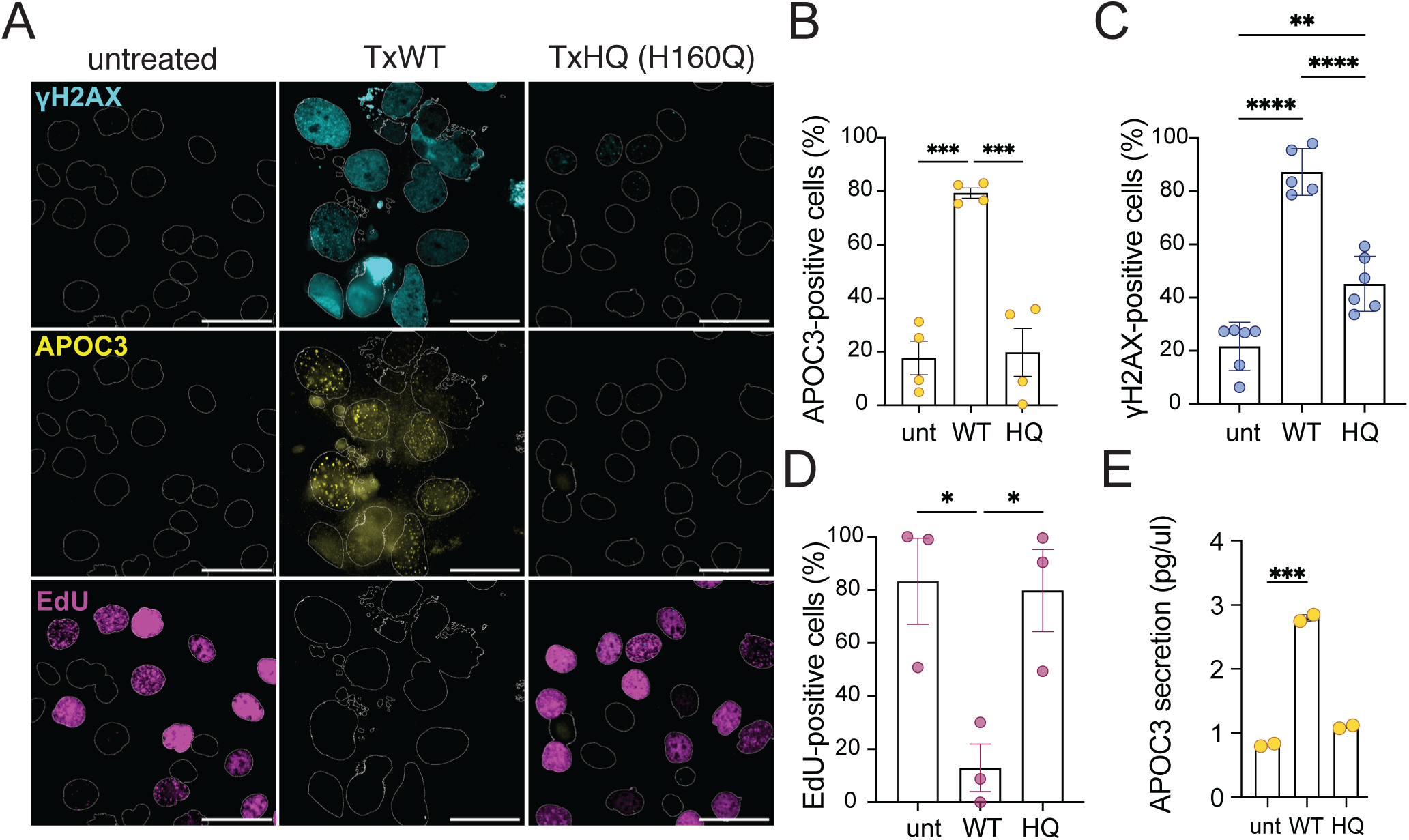
Typhoid toxin-induced DNA damage and APOC3 secretion. **(A)** Representative images of CACO2 intestinal cells either untreated, treated with wild-type typhoid toxin (TxWT) or H160Q DNase-deficient toxin (TxHQ) for 2 hours prior to imaging at 96h of γH2AX (cyan), APOC3 (yellow) and EdU (Magenta). EdU was incubated with cells 24h before fixation. DAPI-stained nuclear outlines shown. Scale bars 50µm. **(B)** Bar chart showing proportion of APOC3 expressing cells. **(C)** Bar chart showing proportion of γH2AX-positive cells. **(D)** Bar chart showing proportion of cells incorporating EdU nucleotide analogue **(E)** ELISA of APOC3 secreted into growth media harvested from cells in (A). Asterisks in **(B), (C), (D)** and **(E)** indicate significance calculated by one way ANOVA, data represented as mean ±SEM. *p<0.05, **p<0.01, ***p<0.001, ****p<0.0001. Circles represent biological repeats.

**Figure 3:**
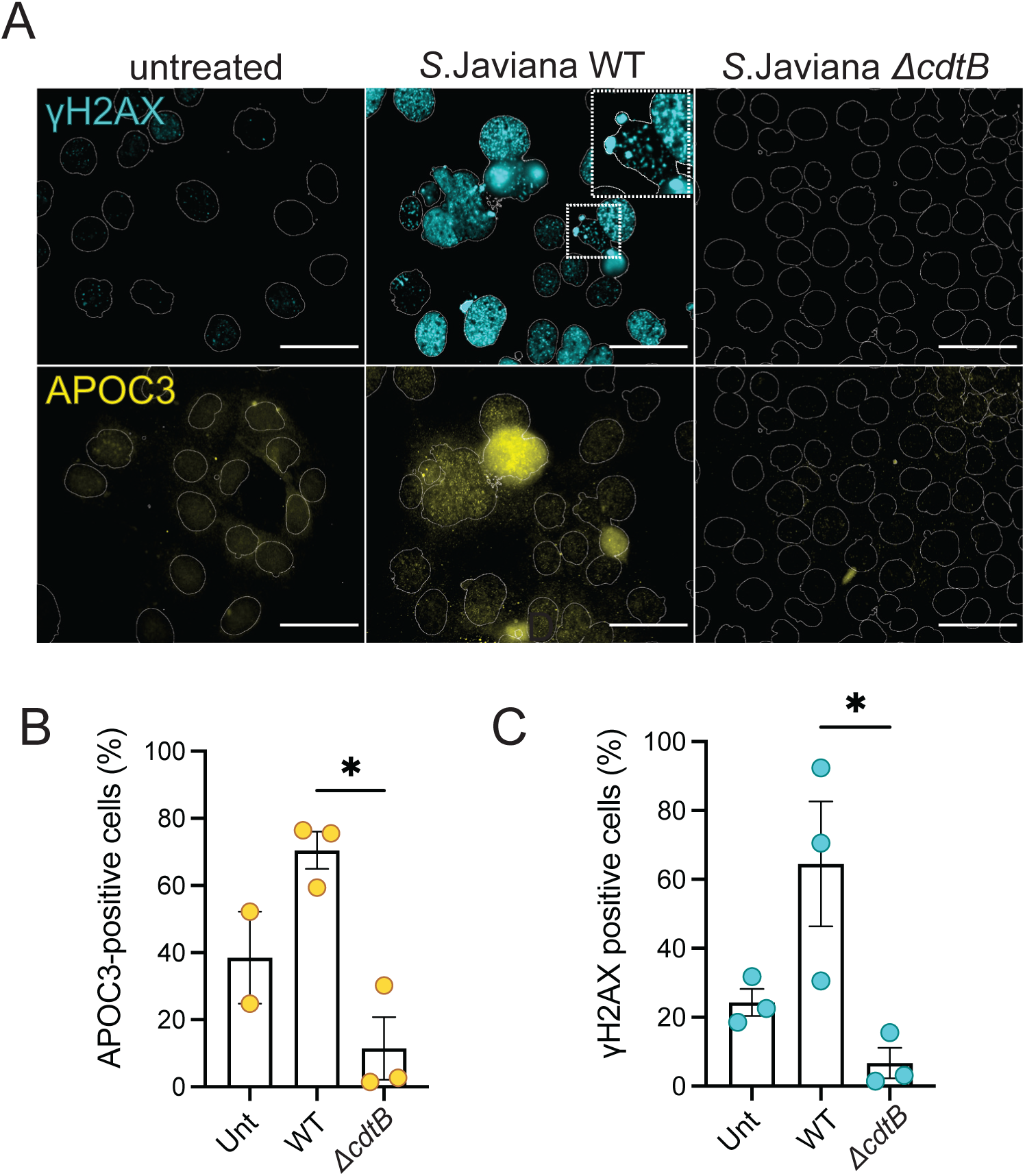
*Salmonella* induces DNA damage and APOC3 expression. **(A)** Representative images of CACO2 intestinal cells infected with wild-type or toxin-deficient (*ΔcdtB*) *S*.Javiana for 1h prior to incubation in gentamicin-containing media and imaging at 96h. Imaging of γH2AX (cyan) and APOC3 (yellow) with DAPI-stained nuclear outlines shown. Magnified inset indicates micronuclei. Scale bars 50µm. **(B)** Bar chart showing proportion of APOC3 expressing cells. **(C)** Bar chart showing proportion of γH2AX-positive cells. Asterisks indicate significance calculated by one way ANOVA, data represented as mean ±SEM. *p<0.05, **p<0.01, ***p<0.001, ****p<0.0001. Circles represent biological repeats.

### Toxin-induced DDRs mediate expression of APOC3 and LYZ

We investigated whether APOC3 treatment of *S.*Javiana influenced pathogen growth but found no effect (**Fig S2A, S2B**). This suggests that APOC3 provides a marker of toxin-induced DDRs rather than playing a direct antimicrobial role against *Salmonella*. Consequently, we turned to LYZ, which has established antimicrobial activities through its ability to break down peptidoglycan in bacterial cell walls and through cationic pore-formation [23]. We determined whether LYZ was, like APOC3, also regulated by toxin-induced DDRs. We examined CACO2 intestinal cells treated with either TxWT or the drug etoposide (ETP) (**Fig 4**), which inhibits topoisomerase causing double stranded DNA breaks [26]. In contrast to untreated control cells, we found that either TxWT or ETP induced γH2AX-labelled DDRs (**Fig 4A**), which was associated with expression of APOC3 and LYZ divergently distributed inside the damaged cells (**Fig 4B, 4C, 4D**). Parallel experiments in THP1 macrophages showed that LYZ was equivalent in TxWT- and TxHQ-treated macrophages (**Fig S3A, S3B**), thus, we continued to investigate LYZ in intestinal epithelial cells. To counteract pathology, the DDR is activated through kinases ATM (ataxia-telangiectasia mutated), which responds to DSBs, and ATR (ATM and rad3-related) that senses single-strand DNA breaks [26]. Both ATM/ATR are inhibited by caffeine [27]. We used caffeine to inhibit ATM/ATR and assess their role in APOC3 and LYZ expression (**Fig 4**). Uniting the functions of ATM and ATR is phosphorylation of their effector γH2AX [26], which was used as a control for DDRs. In the presence of caffeine, the γH2AX response to either TxWT or ETP was suppressed indicating inhibition of ATM/ATR (**Fig 4A, 4E**). Caffeine treatment also disabled the ability of toxin-induced DDRs to drive expression of APOC3 and LYZ, a phenotype also observed with ETP (**Fig 4B, 4C, 4D**). In summary, the results indicate that both APOC3 and LYZ display increased expression in response to ATM/ATR-mediated DDRs by typhoid toxin.

**Figure 4:**
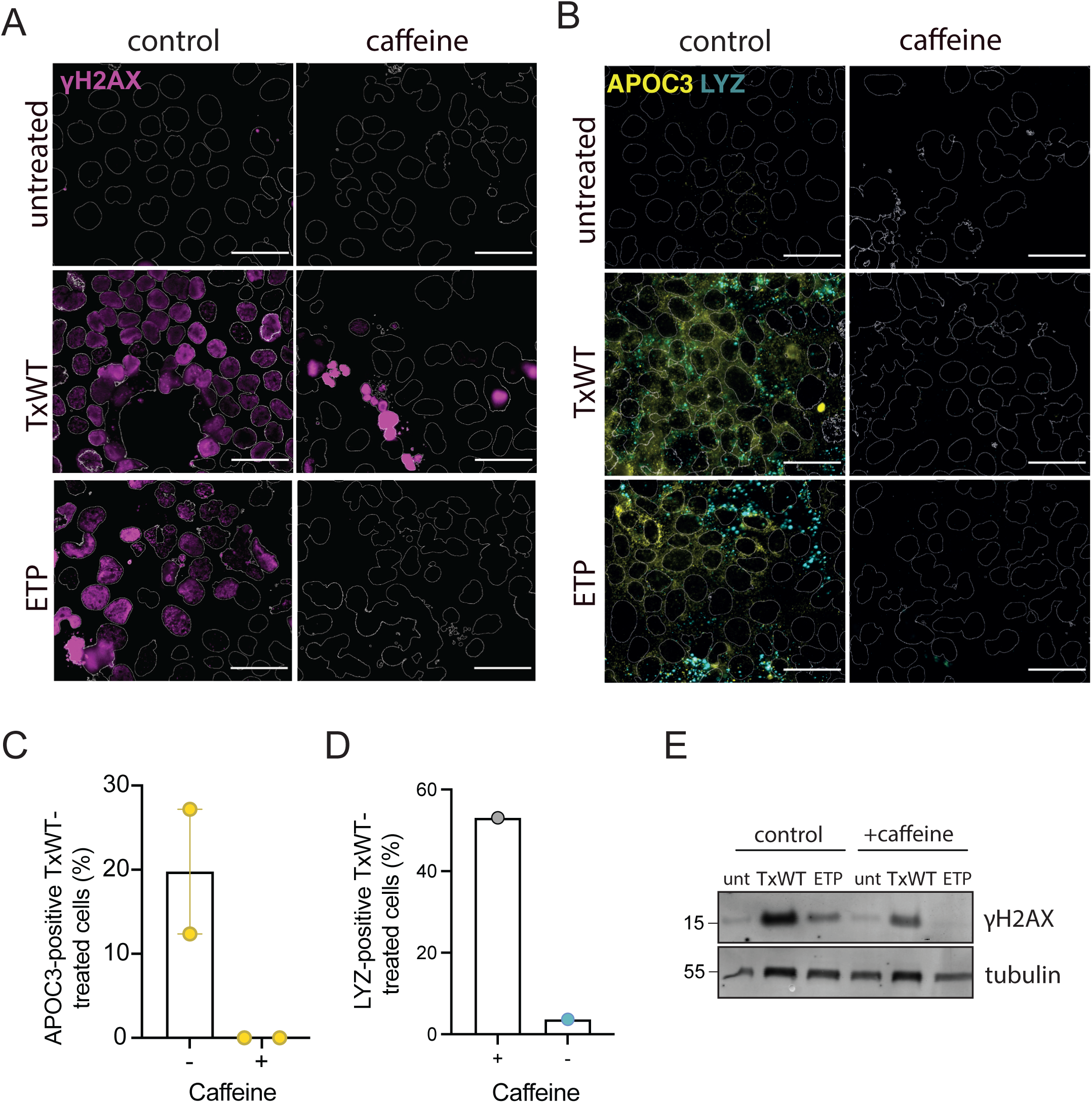
Toxin-induced DNA damage responses mediate APOC3 and LYZ expression. **(A)** Immunofluorescence images of CACO2 cells treated with fresh media (untreated), 20ng/ml TxWT or 8mM etoposide (ETP) for 2 hours with and without continuous caffeine before imaging at 48h. Representative images show γH2AX (magenta) and outlines of DAPI-stained nuclei. Scale bars 50µm. **(B)** Same as (A) with imaging of APOC3 (yellow) and LYZ (cyan). (**C**) Bar chart showing proportion of APOC3-positive cells, or (**C**) LYZ-positive puncta per field of view, in CACO2 cells treated with TxWT in the presence or absence of caffeine. (**C), (D)** Circles represent biological repeats. (**E**) TxWT or ETP-treated cells in the presence or absence of caffeine for 48h immunoblotted with antibodies to γH2AX or the loading control tubulin. MW in kDa, left.

### LYZ causes *Salmonella* spheroplast formation and is augmented by lactoferrin

We investigated LYZ further due to its known antimicrobial properties. Relative to untreated control cells, we found that TxWT induced LYZ expression and an increase in LYZ-positive cells, which was absent in TxHQ-treated cells at 96h, indicating a role for toxin nuclease activity (**Fig 5A, 5B**). The action of TxWT-induced nuclease activity was consistent with γH2AX-labelled DDRs and a lack of EdU incorporation into host cell DNA, showing cell cycle arrest. This contrasted with controls that lacked γH2AX and synthesised EdU-positive DNA (**Fig 5A**: untreated, TxHQ). In addition, we found that only TxWT significantly increased secretion of LYZ relative to controls (**Fig 5C**).

**Figure 5:**
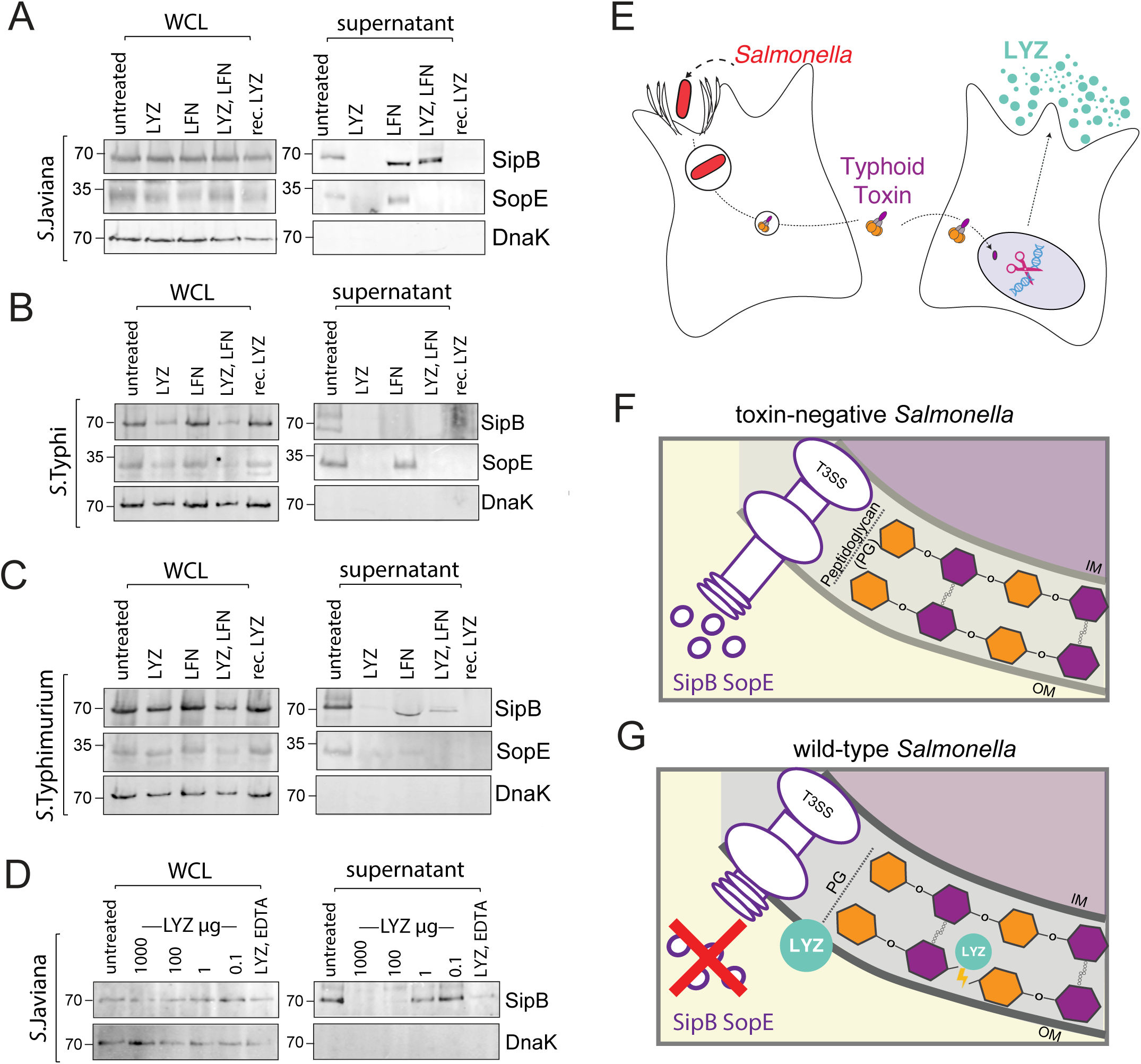
Lysozyme secreted in response to typhoid nuclease activity and causes *Salmonella* spheroplast formation. **(A)** Representative Immunofluorescence images of CACO2 cells either untreated or treated with 20ng/ml TxWT or TxHQ for 2h before imaging at 96h. Images show γH2AX (cyan), LYZ (yellow) and EdU (magenta) with outlines of DAPI-stained nuclei. EdU nucleotide was incubated with cells for 24h prior to fixation at 96h. Magnified inset shows cell cycle-arrested cell producing LYZ. Scale bars 50µm. **(B)** Bar chart showing proportion of LYZ-expressing cells. **(C)** ELISA of LYZ secreted into growth media harvested from cells in (A). **(B), (C)** Circles represent biological repeats. **(D)** Representative transmission electron microscopy (TEM) images of *S.*Javiana either untreated or incubated with 1mg/ml LYZ, 100µg/ml LFN, LYZ and LFN, LYZ and 1mM EDTA in M9 minimal media for 2h. **(E)** Bar chart showing proportion of *S*.Javiana with spherical morphology from the experiment in (D). **(F)** Representative TEM images from the experiment in (D) highlighting changes in cell morphology indicative of *S.*Javiana spheroplast formation in the presence of LYZ and LFN. **(G)** LYZ and LFN treatment of *S*.Javiana in the presence of cephalexin. Left: schematic representation of cell elongation due to cephalexin and subsequent rounding due to spheroplast formation. Right: Microscopy images of *S*.Javiana pFPV-mCherry grown in LB until OD600 1.0 either untreated (left) or treated with cephalexin (right) prior to incubation with cephalexin only (control), LYZ, or LYZ and LFN. Top row: fluorescent images of DAPI-stained (blue) mCherry *S*.Javiana (red) with red arrows indicating elongated cephalexin-treated *S*.Javiana and yellow arrows indicating spheroplasts incapable of mCherry retention due to LYZ and LFN. Bottom row: corresponding phase contrast images. Scale bars 5µm. **(H)** Bar chart showing number of elongated *S*.Javiana (>5µm) per field of view in indicated treatments from experiment in (G). Asterisks in **(E)** indicate significance calculated by one way ANOVA, data represented as mean ±SEM. *p<0.05, **p<0.01, ***p<0.001, ****p<0.0001.

LYZ is best known for its ability to degrade peptidoglycan and induce cationic pore formation [23]. Hydrolysis of peptidoglycan and subsequent spheroplast formation are assisted by factors that mediate LYZ penetration into the periplasm of Gram-negative bacteria [23]. One such factor is lactoferrin (LFN), which destabilises the bacterial cell wall facilitating LYZ entry [28]. LFN was found in all participants by proteomics indicating that LYZ and LFN work in combination to suppress *Salmonella* in response to typhoid toxin. Therefore, we examined whether LYZ and LFN generated spheroplasts. Transmission electron microscopy showed that untreated *S.*Javiana had a rod shaped morphology, or a spherical morphology in ∼50% of cases depending on bacterial cell orientation (**Fig 5D, 5E**). When *S.*Javiana were treated with LFN alone, there was no significant difference and *Salmonella* morphology was equivalent to untreated while LYZ alone induced a small but significant increase in spheroplast formation (**Fig 5D, 5E**). In contrast, LYZ and LFN in combination had a marked effect and increased the proportion of cells with spherical morphology to ∼75%. This was equivalent in significance to the ∼80% of spherical cells observed when LFN was replaced with EDTA that permeabilises the outer membrane of Gram-negative bacteria allowing entry of LYZ. In addition to inducing a round morphology typical of spheroplasts, LFN and LYZ caused instances of membrane damage where cell content lost interaction with its cell membrane resulting in protrusions (**Fig 5F**: magenta arrows). Cell content was also released out of the cell resulting in the formation of a ghost-like shell (**Fig 5F**: blue arrows). We also examined spheroplast formation by fluorescence microscopy using *S.*Javiana expressing mCherry (**Fig 5G**). The change from untreated rod shaped *S.*Javiana to spheroplasts was challenging to observe by fluorescence microscopy due to the <5µm size of bacteria (**Fig 5G**: untreated). To observe spheroplasts more readily, we first treated *S.*Javiana with cephalexin, which inhibits cytokinesis causing an extended morphology (>5µm) that is abolished by spheroplast formation due to degradation of peptidoglycan [29–31] (**Fig 5G**; cartoon left). In the cephalexin-treated control, elongated *S.*Javiana were observed by microscopy that contained mCherry and DAPI-stained DNA (**Fig 5G**; red arrow). In the presence of LYZ and LFN however, the elongated morphology was absent (**Fig 5G**; bottom). Instead, round *Salmonella* that failed to retain mCherry were found by DAPI staining (yellow arrow), which demonstrated spheroplast formation and is consistent with cell content release observed by electron microscopy (**Fig 5F**: LFN, LYZ). The number of elongated *Salmonella* was quantified, which confirmed that LYZ and LFN reduced the number of elongated *Salmonella* (**Fig 5H**). The same trend was observed when LFN was replaced with EDTA. In contrast, LFN alone had no effect as elongated *Salmonella* were still observed (**Fig 5H, S2A**). We also found that LYZ alone caused a loss in cell morphology though the *S.*Javiana, which retained mCherry (**Fig 5G, 5H**). This agrees with observations by electron microscopy where LYZ increased the proportion of spherical cells (**Fig 5D, 5E**) and suggests limited entry of LYZ into the periplasm. The same trends were observed in *S.*Typhi and *S.*Typhimurium where LYZ induced spheroplast formation alone or in combination with LFN or EDTA (**Fig S4B, S4C**).

### LYZ suppresses the function of the *Salmonella* Type 3 Secretion System

*Salmonella* infects cells using a T3SS, which is structured across the inner and outer membranes of the Gram-negative bacterium for injection of virulence effectors into host cells [32]. Thus, we asked whether LYZ and LFN-induced changes in morphology influence T3SS-mediated secretion of virulence effectors SipB and SopE that play key roles in invasion [2]. LYZ and LFN were added to the cultures of toxigenic *S.*Javiana and *S.*Typhi before analysing the expression and secretion of SipB and SopE (**Fig 6A, 6B**). In the untreated control, we found that both *S.*Javiana and *S.*Typhi expressed SipB and SopE. The untreated culture supernatant contained secreted SipB and SopE, which, as expected, lacked the intracellular loading control DnaK that was present in the whole cell lysate (WCL). LYZ and LFN in combination did not impair secretion of SipB in *S.*Javiana but did inhibit SopE (**Fig 6A**). The effect was more striking in *S.*Typhi where secretion of both SipB and SopE was inhibited by LYZ and LFN (**Fig 6B**). The trend observed with *S.*Javiana was also found in *S.*Typhimurium where SopE was predominantly affected (**Fig 6C**). Thus, spheroplast formation impairs the function of the T3SS. In control experiments, we found that LFN alone had no effect on T3SS-mediated secretion in *Salmonella* as secreted SipB and SopE were detected (**Fig 6A, 6B, 6C**). To our surprise however, we found that LYZ alone impaired T3SS as SipB and SopE secretion was lost in *S.*Javiana, *S.*Typhi and *S.*Typhimurium (**Fig 6A, 6B, 6C**). The same trend was observed when recombinant LYZ (rec.LYZ) replaced the endogenous conventional lysozyme (i.e. LYZ). Extracellular LYZ concentrations range from 1200µg/ml in tears to 10µg/ml in the serum of healthy adults [33]. Thus, we examined secretion of SipB by *S.*Javiana treated with indicated concentrations of LYZ (**Fig 6D**). We found that 1000µg/ml and 100µg/ml LYZ abolished *Salmonella* secretion of SipB whose secretion was inhibited, but not abolished, by 1µg/ml LYZ while 0.1µg/ml LYZ was equivalent to the untreated control. It is possible that LYZ-induced pore formation rather than spheroplast formation, caused loss of secretion via the T3SS. However, when we imaged *Salmonella* following cephalexin treatment, we observed that LYZ and rec.LYZ each induced spheroplast formation indicating penetration of lysozyme into the periplasm of *Salmonella* (**Fig 5H**). Taken together, the results indicate that toxin-induced DDRs during typhoid fever induce host secretion of LYZ, which impairs virulence mechanisms of *Salmonella* alone or in combination with LFN.

**Figure 6:**
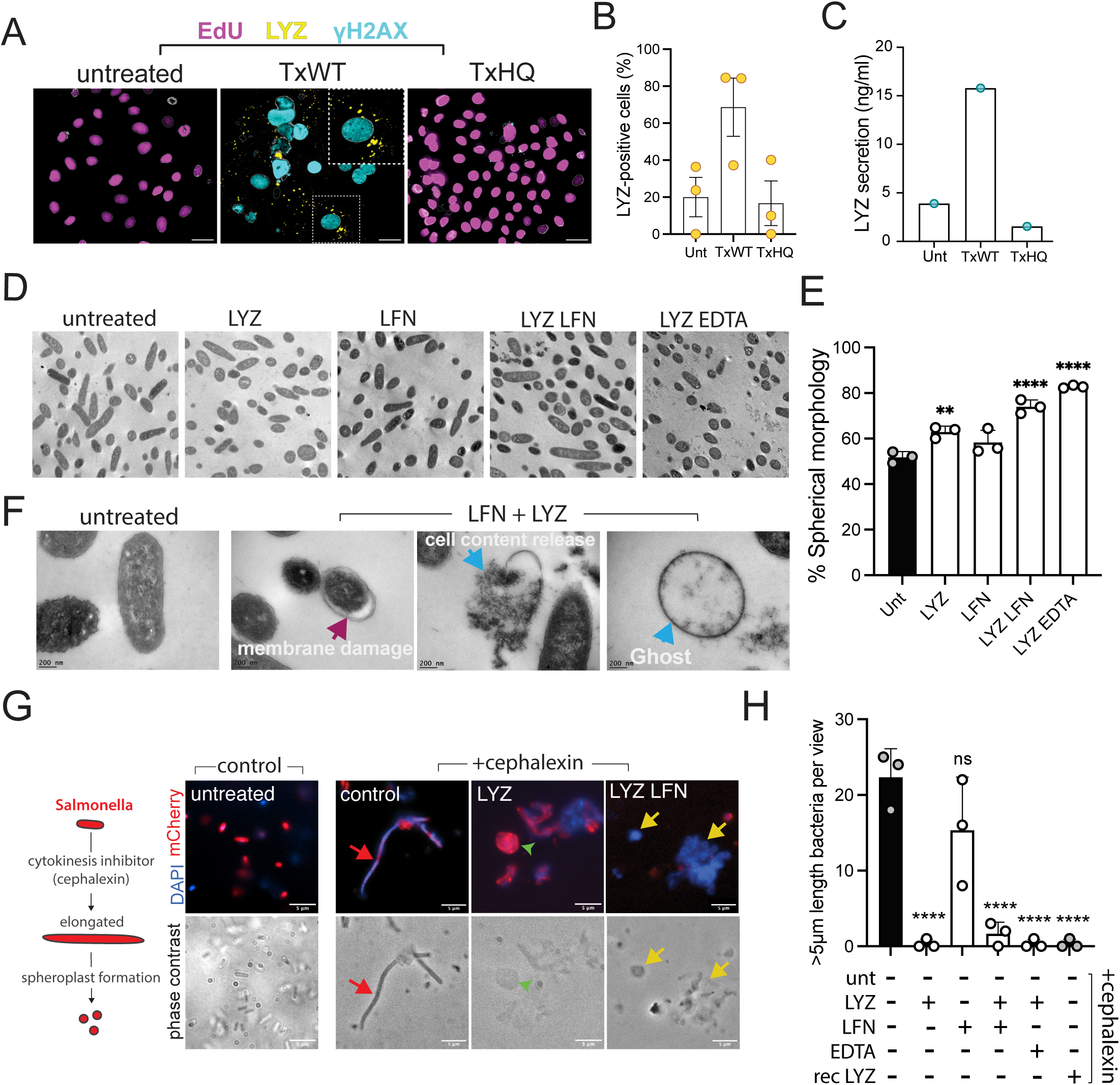
Lysozyme inhibits *Salmonella* T3SS secretion of virulence effector proteins. **(A)** *S.*Javiana, **(B)** *S.*Typhi, or **(C)** *S*.Typhimurium were cultured in LB broth then either left untreated in LB, or cultured with the addition of 1mg/ml endogenous LYZ, 100µg/ml LFN, LYZ and LFN, or 1mg/ml recombinant LYZ (rec. LYZ) for 2h. Whole cell lysates (WCLs) or supernatants containing secreted proteins were harvested for immunoblotting with antibodies to virulence effectors SipB or SopE, or the intracellular loading control DnaK. **(D)** Whole cell lysates or culture supernatants were prepared from *S.*Javiana cultured either untreated in LB or treated for 2h with indicated concentrations of LYZ, or LYZ and EDTA. Samples were immunoblotted with antibodies antibodies to SipB or SopE or DnaK. MW in immunoblots indicated left in kDa. **(E)** Proposed model showing *Salmonella* induction of DNA breaks and consequent host secretion of LYZ. Proposed model showing that **(F)** the type 3 secretion system (T3SS) of toxin-negative *Salmonella* secrete virulence effectors SipB and SopE. In contrast, **(G)** wild-type toxigenic *Salmonella* activate innate defences due to typhoid toxin, which includes host secretion of LYZ that suppresses pathogen secretion of SipB and SopE. LYZ degrades peptidoglycan and causes cationic pore formation in the OM. Inner membrane (IM), outer membrane (OM).

## Discussion

*Salmonella* Typhi and related toxigenic *Salmonella* have evolved a chimeric typhoid toxin implicated in typhoid fever, immune evasion and chronic *Salmonella* carriage [9, 14, 34]. Intravenous injection of purified typhoid toxin causes lethality in a mouse model, which is dependent on toxin nuclease activity [9, 34, 35]. This prompted a human infection challenge study comparing wild-type and toxin-negative *S.*Typhi [16], which showed that participants infected with wild-type *S.*Typhi had significantly shorter duration of bacteraemia, a hallmark of typhoid fever. Thus, we hypothesised that toxin-induced DDRs may induce an innate immune response that suppresses infection. Our study demonstrates that participants infected with wild-type *S.*Typhi induce a divergent host secretome, which contains effectors of innate immunity including the antimicrobial lysozyme (**Fig 6E: Proposed model**). Indeed, we observed that toxin-mediated DDRs triggered secretion of lysozyme in cultured cells and that lysozyme inhibited the function of the *Salmonella* T3SS (**Fig 6F, 6G**). This is the first time typhoid toxin has been associated with eliciting host defence mechanisms that protect humans from typhoid fever.

The findings from this study support the view that the activity of typhoid toxin comes at a cost for *Salmonella* during typhoid fever, which leaves the role of the typhoid toxin unclear. Indeed, *S.*Paratyphi B does not encode typhoid toxin but can cause a typhoid-like disease [36], which was observed with *S.*Typhi-TN [16]. It is possible that the presence of typhoid toxin in the blood causes fatalities, which has been observed in mouse models [9, 34, 35], and is a symptom associated with severe typhoid fever in humans [1]. However, human infection challenge studies cannot test severe typhoid fever for ethical reasons [16]. It is also possible that the cost of activating host defences is balanced somewhat by the role of typhoid toxin in persistent infections and systemic spread, which has been observed during mouse infection models [14, 15] Indeed, this study further highlights the arms race between host and pathogen by providing an example of how infection is counteracted during typhoid fever.

The typhoid toxin induces cellular senescence through DNA replication stress and mitochondrial injury in cancer cells, primary macrophages and lymphocytes [10, 13, 37], and in mouse infection models [11, 37]. Senescence is typically activated in response to DDRs and is best known as a tumour suppressor mechanism but may have co-evolved to combat bacterial infections [38, 39]. Host secretomes of senescent cells underlying the senescence-associated secretory phenotype (SASP) are a likely source of antimicrobial activity. Indeed, IFNγ, IFN-1, TNFα, IL-1β, and CCN1 have been implicated in antimicrobial responses [40–44], and each have been identified as SASP factors from diverse senescence triggers, e.g. DNA replication, wound healing and oncogene activation [39]. Though SASP has not been investigated for its antimicrobial properties, we propose that lysozyme fits with the type of diverse type of antimicrobial factors already identified as SASP factors.

Lysozyme degrades peptidoglycan whose activity in Gram-negative bacteria is augmented by lactoferrin, which was identified in participants by proteomics. Lysozyme and lactoferrin have previously been shown to inhibit the growth of *S.*Typhimurium [28]. In support of this, we found that *Salmonella* treated with lysozyme and lactoferrin failed to retain fluorescent proteins, which discriminates live from dead *Salmonella* [45, 46]. Consistent with effects on protein retention, lysozyme has also been shown to liberate peptidoglycan-binding protein Lpp1 from the outer membrane of *S.*Typhimurium in infected mice causing pro-inflammatory responses [47]. We found that lysozyme and lactoferrin combined to generate spheroplasts that suppressed T3SS-mediated secretion of SopE. However, the inhibitory effects on T3SS activity was greater with lysozyme alone. As a result, we cannot exclude LYZ functions that occur independently of lactoferrin such as LYZ-mediated membrane pore-forming activity, which may play a role in inhibiting T3SS.

Host secretomes undergoing toxin-induced DNA damage were identified by proteomics in the absence of serum [12] that contains abundant plasma proteins, which mask less abundant proteins during proteomics [21]. Though we depleted the most abundant plasma proteins in infection challenge samples, our proteomics identified relatively abundant proteins such as APOC3 and lysozyme. This is despite identification of cytokines in the TYGER challenge study investigating typhoid toxin [16], which indicates there are additional unidentified host stress mechanisms to be discovered that suppress the duration of bacteraemia in response to typhoid toxin. Indeed, *Salmonella* is known to encode inhibitors of lysozyme [23] and we think it unlikely that lysozyme acts alone in counteracting *Salmonella* following DNA damage by the typhoid toxin. This study presents the view that pathogen induction of host DNA damage responses elicits antimicrobial responses, which impact infectious disease and explain the shorter duration of bacteraemia in participants infected with wild-type toxigenic *S.*Typhi.

## Materials and Methods

### Immunodepletion of abundant plasma proteins

Human blood plasma was obtained with ethical approval from the South-Central Oxford A Ethics Committee in the project entitled ‘Investigating Typhoid Fever Pathogenesis (TYGER)’ (Ref: 16/SC/0358), clinicaltrials.gov reference NCT03067961 [16]. Plasma was standardised with respect to protein concentrations using the Micro BCA Protein Assay Kit (Thermo Scientific™, #23235). To immunodeplete the 14 most abundant proteins (albumin, IgG, antitrypsin, IgA, transferrin, haptoglobin, fibrinogen, α2-macroglobulin, α1-acid glycoprotein, IgM, apolipoprotein AI, apolipoprotein AII, complement C3 and transthyretin), plasma (1.5µg/µl) was diluted 20-fold in Buffer A (proprietary buffer from Agilent; #5188-6559) prior to centrifugation and removal of any particulates (5 mins, 16,000xg). Diluted plasma was applied to the Multiple Affinity Removal Column Human-14 (MARS-14, Agilent, #5188-6559). Low-abundant proteins were collected in the flow-through and high-abundant proteins remained bound to MARS-14 and were eluted in Buffer B (proprietary buffer from Agilent, #5188-6559). Eluted low-abundant proteins were precipitated using acetone to concentrate proteins prior to mass spectrometry analysis. Acetone was added to low-abundant protein elution at −20°C (4:1 ratio). Precipitated proteins harvested by centrifugation (10 mins, 16,000xg at 4°C). Air-dried pellets resuspended in 50µl of S-Trap solubilization buffer (5% SDS, #05030-500ML-F; 5mM Triethylammonium Bicarbonate Buffer (TEAB); #90114;, pH 7.55) and stored at −20°C ready for further analysis.

### Sample preparation for mass spectrometry analysis

Samples in S-Trap solubilization buffer were reduced with TCEP (#646547-10X1ML, Sigma-Aldrich) at a final concentration of 10 mM and heated to 70°C for 15 minutes. Following this, the samples were incubated with iodoacetamide (#I6125, Sigma-Aldrich) at a final concentration of 10 mM and stored in the dark at room temperature. Phosphoric acid (#A242-500, Sigma-Aldrich) was added to a final concentration of 1.2%. The samples were then diluted in S-Trap binding buffer (90% aqueous methanol, #900688-1L, Sigma-Aldrich; 0.1M TEAB, pH 7.1). The diluted samples were loaded onto S-Trap columns (#C02-micro-80, Protifi) by centrifugation (10 seconds at 4,000 × g, room temperature) and digested with trypsin. Trypsin (#90058, Thermo Fisher Scientific) in 0.1% trifluoroacetic acid (TFA) (#108262, Sigma-Aldrich) was added at a ratio of 1 µg trypsin per 10 µg of protein immobilized on the S-Trap columns. The S-Traps were sealed with parafilm and incubated at 47°C for 1 hour. Digested proteins were eluted with 50 mM TEAB, followed by centrifugation at 4,000 × g for 10 seconds. Aqueous formic acid was added to the eluate to a final concentration of 0.1% after initially adding 0.2%, and the mixture was centrifuged at 4,000 × g for 10 seconds. Subsequently, 40% of 50% acetonitrile containing 0.2% aqueous formic acid (#A117-50, Sigma-Aldrich) was added, and the samples were centrifuged again under the same conditions.The samples were stored at −20°C until they were dried. For drying, they were centrifuged at 45°C for 90 minutes in a vacuum at 1,400 rpm using an Eppendorf™ Concentrator Plus. Once dried, the samples were resuspended in 0.5% formic acid, transferred to polypropylene vials (Thermo Scientific, #160134), and injected into the Orbitrap for LC-MS/MS analysis.

### LC-MS/MS analysis of proteomic data

Samples were analysed by nanoflow LC-MS/MS using an Orbitrap Elite (Thermo Fisher) hybrid mass spectrometer equipped with an easyspray source, coupled to an Ultimate RSLCnano LC System (Dionex). Xcalibur 3.0.63 (Thermo Fisher) and DCMSLink (Dionex) controlled the system. Peptides were desalted on-line using an Acclaim PepMap 100 C18 nano/capillary BioLC, 100A nanoViper 20 mm x 75 µm I.D. particle size 3 µm (Fisher Scientific) followed by separation using a 125-min gradient from 5 to 35% buffer B (0.5% formic acid in 80% acetonitrile) on an EASY-Spray column, 50 cm Å∼ 50 µm ID, PepMap C18, 2 µm particles, 100 °A pore size (Fisher Scientific). The Orbitrap Elite was operated with a cycle of one MS (in the Orbitrap) acquired at a resolution of 60,000 at m/z 400, with the top 20 most abundant multiply charged (2+ and higher) ions in a given chromatographic window subjected to MS/MS fragmentation in the linear ion trap. An FTMS target value of 1e6 and an ion trap MSn target value of 1e4 were used with the lock mass (445.120025) enabled. Maximum FTMS scan accumulation time of 500 ms and maximum ion trap MSn scan accumulation time of 100 ms were used. Dynamic exclusion was enabled with a repeat duration of 45 s with an exclusion list of 500 and an exclusion duration of 30 s.

### MaxQuant Analysis of proteomic data

All raw mass spectrometry data were analysed with MaxQuant version 1.6.10.43. Data were searched against a human UniProt sequence database (May 2019) using the following search parameters: digestion set to Trypsin/P with a maximum of 2 missed cleavages, methionine oxidation and N-terminal protein acetylation as variable modifications, cysteine carbamidomethylation as a fixed modification, match between runs enabled with a match time window of 0.7 min and a 20-min alignment time window, label-free quantification enabled with a minimum ratio count of 2, minimum number of neighbours of 3 and an average number of neighbours of 6. A first search precursor tolerance of 20ppm and a main search precursor tolerance of 4.5 ppm was used for FTMS scans and a 0.5 Da tolerance for ITMS scans. A protein FDR of 0.01 and a peptide FDR of 0.01 were used for identification level cut-offs.

### Perseus Bioinformatic Analysis of proteomic data

MaxQuant data output was loaded into Perseus version 1.6.10.50 and all LFQ intensities were set as main columns. The Matrix was filtered removing any contaminants, identified by site and reverse sequences. LFQ intensities were transformed using Log2(x) default function. Rows were then filtered with a minimum value of 5 in each group only showing the default valid values. Data was visualised using Pearson correlation values and outliers omitted. Sample 8183 D0 was excluded from the analysis due to an inconsistent Pearson correlation value. Missing values were then replaced with a width of 0.3 and down shift 1.8. Sample groups where then compared two at a time with D0 and TD of respective groups using student t-test with permutation-based FDR calculation (FDR = 0.05) with an S0=0.1. Data was then exported to Microsoft excel and Graphpad prism before figure assembly. T h e m a s s s p e c t r o m e t r y p r o t e o m i c s data have been deposited to the ProteomeXchange Consortium via the PRIDE partner repository with the dataset identifier PXD058381.

### Cell Culture

CACO-2 (ATCC #HBT-37), HepG2 (ATCC #HB-8065) and THP1 (ATCC #TIB-202) cells were stored in cryopreservative media (10% DMSO (Sigma-Aldrich, #D2438) and 90% complete media as below at −80°C until being revived in culture. Frozen cells were thawed at 37°C for 90 secs and cultured in recommended media. CACO-2 in complete medium containing Gibco Dulbecco modified eagle media (ThermoFisher, #31966021) with 10% FBS, 1% Penicillin/Streptomycin (#11548876) and 50µg/ml Kanamycin. HepG2 cells were cultured in MEM (Merck #M0518) with 10% FBS, 1% Penicillin/Streptomycin (#11548876) and 50µg/ml Kanamycin. Cells were cultured at 37°C humidified incubator with 5% CO2. Cells were passaged every 3 to 5 days depending on their doubling time. For sub-culturing cells, trypsin (Sigma Aldrich, #T4049) was used to detach adherent cells followed by neutralisation with FBS (Sigma Aldrich, #F7524) containing media. Cell viability was determined before splitting using trypan-blue (Sigma-Aldrich, #T8154) in 1:1 dilution, followed by cell counting of viable trypan-blue negative cells and non-viable trypan blue-positive cells using a hemocytometer (Hawksley, #AC1000). Before seeding, glass coverslips (VWR, #631-1578) were added into cell culture wells and cells diluted in complete growth media and cultured at 37°C humidified incubator with 5% CO2. If necessary, conditioned media was harvested by centrifugation at 6000 × g for 5 mins to pellet the cells before filtering through 0.2µm filters to remove any floating cell debris and storage at −80°C ready for use. To assay for cell cycle arrest, Click-iT^TM^ EdU Cell Proliferation Kit for Imaging, Alexa FluorTM 647 dye (Thermofisher, C10340) was used per the manufacturer’s instruction. EdU was added to the culture 24h before fixation.

THP1 cells were cultured in RPMI-1640 (ThermoFischer #R8758-500ML) in 10%FBS, 1% Penicillin/Streptomycin and 50µg/ml Kanamycin at a cell suspension density of 2-10×10^5^ cells per ml. THP1 differentiation to M1 macrophages was carried out by incubating THP1 undifferentiated monocytes in 10ng/ml Phorbol 12-myristate 13-acetate (PMA, Merck, #P1585) for 48 hours. PMA was washed off thrice in PBS before incubation in complete media containing 50ng/ml IFN-γ (Merck Millipore, #IF002) and 15ng/ml lipopolysaccharide (Thermofisher, #L23352) for 24 hours. During intoxication assays, IFN-γ and LPS were removed.

### Recombinant toxin purification and intoxication assays

The typhoid toxin was purified from BL21 DE3 pETDuet-1 encoding *pltB*^His^ *pltA*^Myc^ and *cdtB*^FLAG^ using NiNTA agarose (Qiagen) affinity chromatography according to manufacturer instructions as previously described (Ibler et al 2019). Unless stated otherwise, cells were intoxicated with 20 ng/ml toxin (∼175 picomolar) for 2h, washed three times with sterile PBS (Sigma-Aldrich, #J60801.K3) to remove any extracellular toxin and chase with fresh complete growth media for the duration of the experiment.

#### Salmonella

Serovars of *Salmonella enterica* in the study were maintained on LB agar plates and cultured in LB broth. *S.*Javiana (S5-0395) and Δ*cdtB* (M8-0540) [15] were kind gifts from Prof. Martin Weidmann (New York). Vaccine candidate *S*.Typhi Ty2 BRD948 *ΔaroA ΔaroC ΔhtrA* [48] was a kind gift from Prof. Gordon Dougan (Cambridge). *S.*Typhimurium SL1344 was a kind gift from Prof. Vassilis Koronakis (Cambridge).

### *Salmonella* infection

*S.*Javiana wild-type or Δ*cdtB* encoding pM975, which expresses GFP when bacteria are intracellular(57) were cultured in LB 50µg/ml ampicillin at 37°C in a shaking incubator to 2.0 OD600. The multiplicity of infection (MOI) was optimised for CACO2 cells (MOI 100). To assay *Salmonella-*induced host cell signalling, infection was initiated in the absence of antibiotics by addition of *Salmonella* to cell cultures in complete growth medium and centrifugation for 1 min at 1000 × g followed by 30 min incubation at 37°C 5% CO_2_. Infected cells were washed three times with PBS and incubated in growth media containing 50 µg/ ml gentamicin (Chem Cruz, sc203334) for 1.5h then reduced to 10 µg/ml gentamicin for the rest of the experiment. When assaying *Salmonella* invasion, the method was modified by serum-starving cells 24h prior to infection that deprives cells of membrane ruffling stimulants in FBS. To assess invasion, *Salmonella* infection occurred over 30 mins. After 1.5h incubation with 50 µg/ml gentamicin, cells were washed three times with PBS and lysed with 1% Triton X-100. Serial dilutions of cell lysates were used to inoculate (5 µl) LB agar plates containing 50µg/ml ampicillin and the *Salmonella* cultured overnight at 37°C. *Salmonella* colony counts were used to quantify colony forming units (CFUs).

### *Salmonella* treatment with APOC3 and Lysozyme

Endogenous conventional LYZ (Sigma-Aldrich, #L2879-1G) or recombinant human conventional LYZ (Sigma-Aldrich, #L1667) were resuspended in 10mM Tris-HCl pH8 at 10mg/ml. *Salmonella* was grown in either LB or M9 minimal media before addition of LYZ at concentrations spanning 0.1µg/ml to 1000µg/ml as indicated accordingly in figure legends. Purified APOC3 (Novus Biologics; NBP1-99294) at 50mg/ml were diluted with bacteria in M9 minimal media.

### Immunoblotting

For immunoblotting *Salmonella*, bacteria were cultured from 0.2 OD600 in LB broth in the presence or absence of LYZ (0.1-1000µg/ml), recombinant LYZ (1000µg/ml), LFN (100µg/ ml) or EDTA (1mM) for 2h. To generate whole cell lysates, *Salmonella* were harvested at 4000 RCF for 15 minutes and resuspended in SDS-UREA (50mM Tris-HCl pH 6.8, 2%

SDS, 6M UREA, 0.3% Bromophenol Blue) containing 5% β-Mercaptoethanol (Sigma-Aldrich, #M6250). To harvest supernatants, culture supernatant was filter sterilised using 0.45µm filters (Sigma-Aldrich, #SLHAR33SS) before adding 10% v/v trichloroacetic acid (TCA from Sigma-Aldrich, #91228) to precipitate proteins overnight at 4°C. Precipitated proteins were harvested by centrifugation at 10,000 RCF, 30 minutes at 4°C. Supernatant was discarded and precipitated proteins washed in 100% acetone by centrifugation at 10,000 RCF, 30 minutes at 4°C. Sample was then left to air-dry and precipitated proteins resuspended in SDS-UREA in a volume according to the OD 600 reading of cultures at the 2h point. For immunoblotting mammalian cells, whole cell lysates were generated by re-suspending cultured cells in SDS-UREA. Proteins were separated by 12% Bis-Tris SDS-PAGE gels in MOPS buffer (50 mM MOPS, 50 mM Tris, 0.1% SDS, 20 mM EDTA) and transferred to PVDF transfer stacks (#1704274, Bio-Rad) using Trans-Blot Turbo Transfer System (Bio-Rad). PVDF membranes were blocked with TBS pH7.4 5% non-fat dried milk with antibody incubations and washes performed in TBS pH7.4 0.1% Tween® 20. IRDye-labelled secondary antibodies were used according to manufacturer’s instructions and immunoblots imaged using Odyssey Sa (LiCor).

### Antibodies for immunoblotting and immunofluorescence

Antibodies were purchased from Millipore (yH2AX diluted 1:1000, #05-636-I), GeneTex (APOC3, diluted 1:250, #GTX129994), ThermoFisher, (LYZ, diluted 1:250, #MA1-82873) and Abcam (tubulin diluted 1:1000, #ab61065). Secondary antibodies were diluted 1:10000 and purchased from ThermoFisher Scientific (Alexa 488 donkey anti-mouse IgG, #A-21202; Alexa 594 donkey anti-rabbit IgG, #A-21207) for immunofluorescence microscopy and LiCor Biosciences (IRDye® 800CW Donkey anti-Mouse IgG, 9#25-32212; IRDye® 680RD Donkey anti-Rabbit IgG, 9#26-68073) for immunoblotting.

### Cephalexin treatment

*S.*Javiana pFPV-mCherry [49], *S.*Typhi BRD948 or *S.*Typhimurium SL1344 were cultured in LB broth until an OD600 of 0.7. Bacteria were diluted 1:10 and grown in LB broth with 50µg/ ml Cephalexin for 2.5-3h at 42°C, 120rpm. Culture was diluted 1:10 in LB Broth supplemented with 0.8mM IPTG for 1hr at 37°C, 120rpm. Bacteria were harvested by centrifugation and resuspended in 1M sucrose on ice. 1ml *Salmonella* suspensions were incubated with 80µl of Tris-HCL pH8 with or without addition of LYZ (1mg/ml), EDTA (1mM) or LFN (100µg/ml) at room temperature for 20 minutes. Samples were fixed using PBS 4% paraformaldehyde and air-dried on glass coverslips ready for imaging by microscopy.

### Immunofluorescence microscopy

At experimental endpoints, CACO2 cells were washed with PBS (Biotech, PD8117) then fixed with PBS 4% paraformaldehyde (PFA) (ThermoFisher, #J61899) for 10-15 mins at room temperature. Cells were washed two more times with PBS then blocked and permeabilised using a 3% BSA (Sigma-Aldrich, 1073508600), 0.2% Triton X-100 (VWR, 28817.295) in PBS at room temperature for 1h. Primary and secondary antibodies were incubated with cells in blocking buffer consecutively for 1h and 30min, respectively, then washed with PBS then water and left to dry. Coverslips were then mounted and counterstained on 6µl of VectaShield mounting agent with DAPI (Vector Lab, H1200), and sealed with nail varnish before being imaged on Nikon’s Inverted Ti eclipse equipped with an Andor Zyla sCMOS camera (2560 x 2160; 6.5µm pixels). The objectives used were Plan Apo 10x (NA 0.45); Plan Apo 20x (NA 0.75); Plan Fluor 40x oil (NA 1.3); Apo 60x oil (NA 1.4); Plan Apo 100x Ph oil (Na 1.45); Plan Apo VC 100x oil (NA 1.4). Quad emission filters for used with SpectraX LED excitation (395nm, 470nm, 561nm, 640nm). The imaging software used was NIS elements software.

### Electron Microscopy

Specimens were fixed overnight in a solution of 2.5% glutaldehyde in 0.1M sodium cacodylate buffer. The following day excess fixative was removed and washed free of excess fixative in two changes of 10 minutes each in 0.1M sodium cacodylate buffer. Samples were then post-fixed 2% aqueous osmium tetroxide for 2h. Samples were washed free of excess Osmium tetroxide in buffer solution and dehydrated through a graded series of ethanol solutions in water (50%, 75%, 95%, 100% twice and dried 100% ethanol), and cleared of excess ethanol in epoxypropane (EPP) and then infiltrated in a 50/50 araldite resin: EPP mixture overnight on a rotor. The next day this mixture was replaced with two changes over 8h of fresh araldite resin mixture before being embedded into EM moulds in fresh araldite resin and cured in a 60°C oven for 48-72h. Ultrathin sections, approximately 85nm thick, were cut using a Diatome histoknife on a Reichart Jung ultracut E ultramicrotome and picked up onto formvar coated 200 mesh copper grids. These were stained for 30 mins with saturated aqueous uranyl acetate, blotted free of excess stain using filter paper and washed in distilled water. Sections were counterstained in Reynold’s lead citrate, washed in water again and blotted dry. Sections were examined using an FEI Tecnai Spirit Biotwin 120 Transmission Electron Microscope at an accelerating voltage of 80Kv. Electron micrographs were recorded using Gatan Orius 1000 digital camera and Gatan Digital Micrograph software.

### ELISA

APOC3 (ThermoFisher, #EHAPOC3) and LYZ ELISA kits (Abcam, ab108880) were used as per manufacturer’s instructions. In brief, conditioned media was collected from cultured cells, centrifuged at 6000 RCF for 5min to pellet bacterial cells and the supernatant passed through 0.2µm filters. Filtrate was serially diluted with a factor of 5 to ensure efficient sensitivity before incubation in pre-coated ELISA plates overnight at 4°C. The supernatant was removed and bound proteins washed using proprietary buffer before addition of biotin -labelled primary antibody conjugate to APOC3 or LYZ for 1h at room temperature. Streptavidin-HRP (horseradish peroxidase) was incubated for 1h at room temperature followed by tetramethylbenzidine substrate of HRP for 30mins. Plates were imaged on FLUOstar Omega at 450nm absorbance and a standard curve was generated, sample concentration was extrapolated from standard curve and normalised to cell density.

### Drugs

Drugs were resuspended as per manufacturer’s instructions and diluted in media to their working concentrations. Etoposide was used at 10µM, Aphidicolin was used at 20µM and caffeine at 10mM. CACO2 cells in complete media were incubated with caffeine at 10mM for 30 minutes before intoxication with purified typhoid toxin for 2 hours. Media was replaced with fresh media containing 10mM caffeine and incubated for 48 hours at 37°C in a humidified incubator with 5% CO2.

### Image processing

For qualitative purposes images were processed using fiji 2.0.0-rc-69/1.52p, with a macro code to automate image processing. The macro code adjusts brightness and contrast and is normalised across all images. DAPI nuclei are outlined and overlapped with other channels and the images are then saved as a png file. For the Cell Profiler image quantification, Cell profiler pipeline was used to identify nuclei and quantify the intensity of signals per nuclei. Further pipelines were adjusted from the original pipeline to cater for more experiments.

### Statistical Analysis

Statistical analysis was carried out by Graphpad Prism 9 version 9.0.2. Most analysis are t-test or ANOVA as indicated in figure legends. The significance is represented using *, p < 0.05 (*), p < 0.01(**), p < 0.001 (***), and p < 0.0001(****).

## Figures

All figures were assembled on Adobe Illustrator version 25.4.1.

---

**Figure S1:**
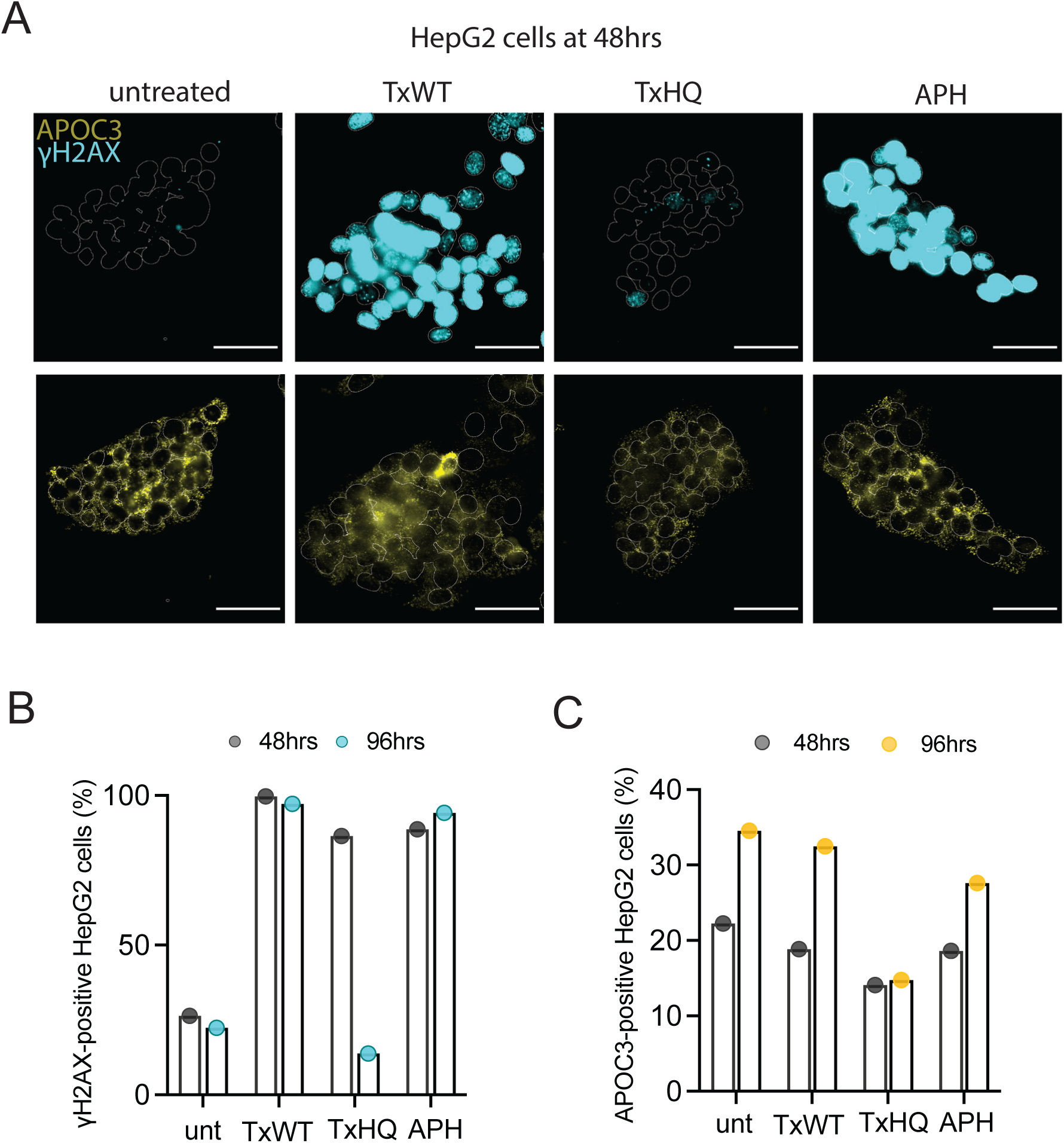
APOC3 expression in HepG2 liver cells treated with typhoid toxin. **(A)** Representative images of HepG2 intestinal cells either untreated, treated with wild-type typhoid toxin (TxWT) or H160Q DNase-deficient toxin (TxHQ) for 2 hours prior to imaging at 48h of γH2AX (cyan) or APOC3 (yellow). DAPI-stained nuclear outlines shown. Aphidicolin was used as a positive control for induction of DDRs and was incubated for the duration of the experiment. Scale bars 50µm. **(B)** Bar chart showing proportion of γH2AX-positive cells at 48h and 96h. **(C)** Bar chart showing proportion of APOC3 expressing cells at 48h and 96h. Circles indicate number of biological repeats.

**Figure S2:**
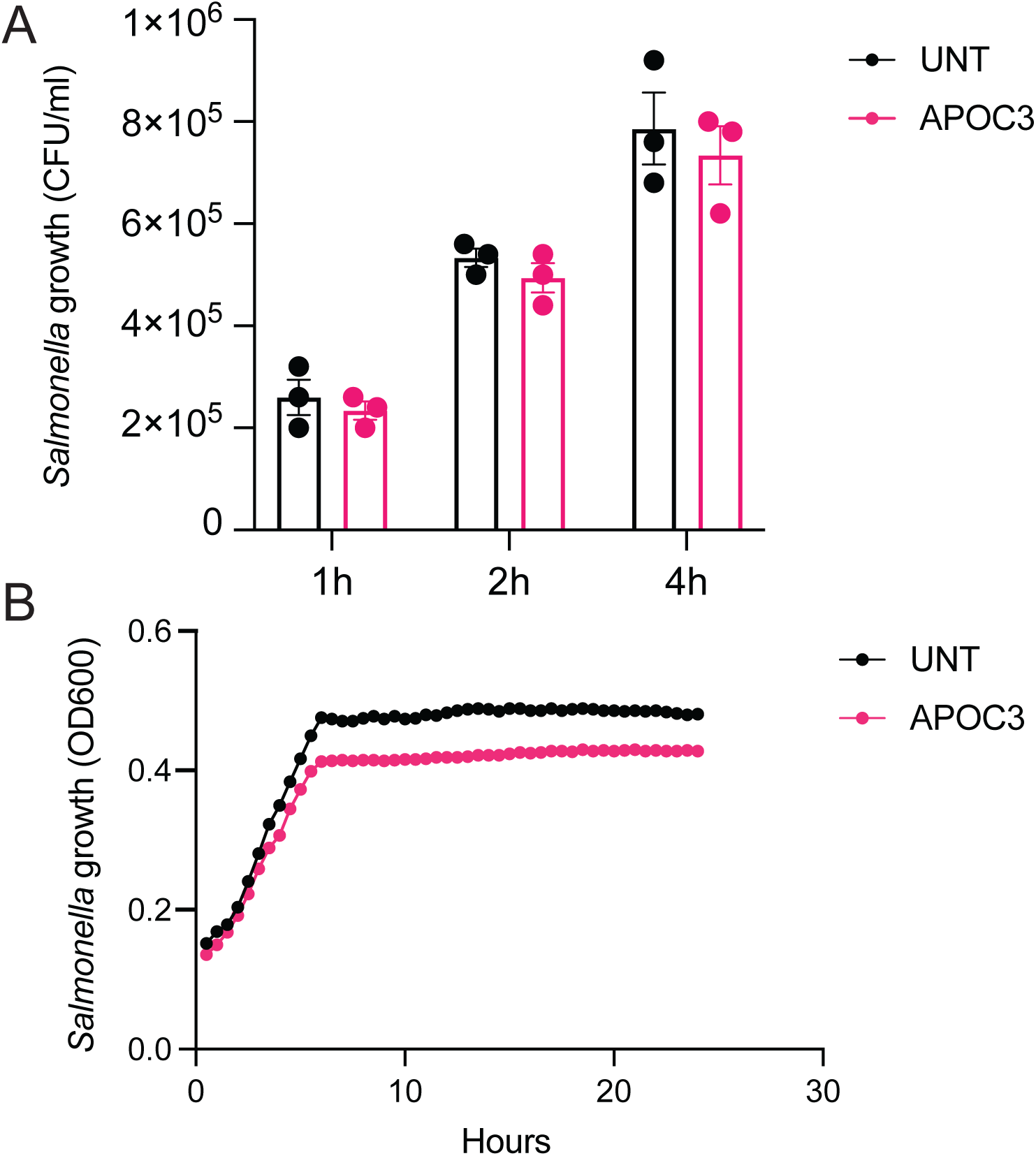
APOC3 has no effect on *Salmonella* growth. **(A)** Quantification of *Salmonella* colony forming units (CFU) following incubation with 50mg/ml of purified APOC3 at 1h, 2h and 4h. **(B)** The same experiment as (A) over a 24hr period.

**Figure S3:**
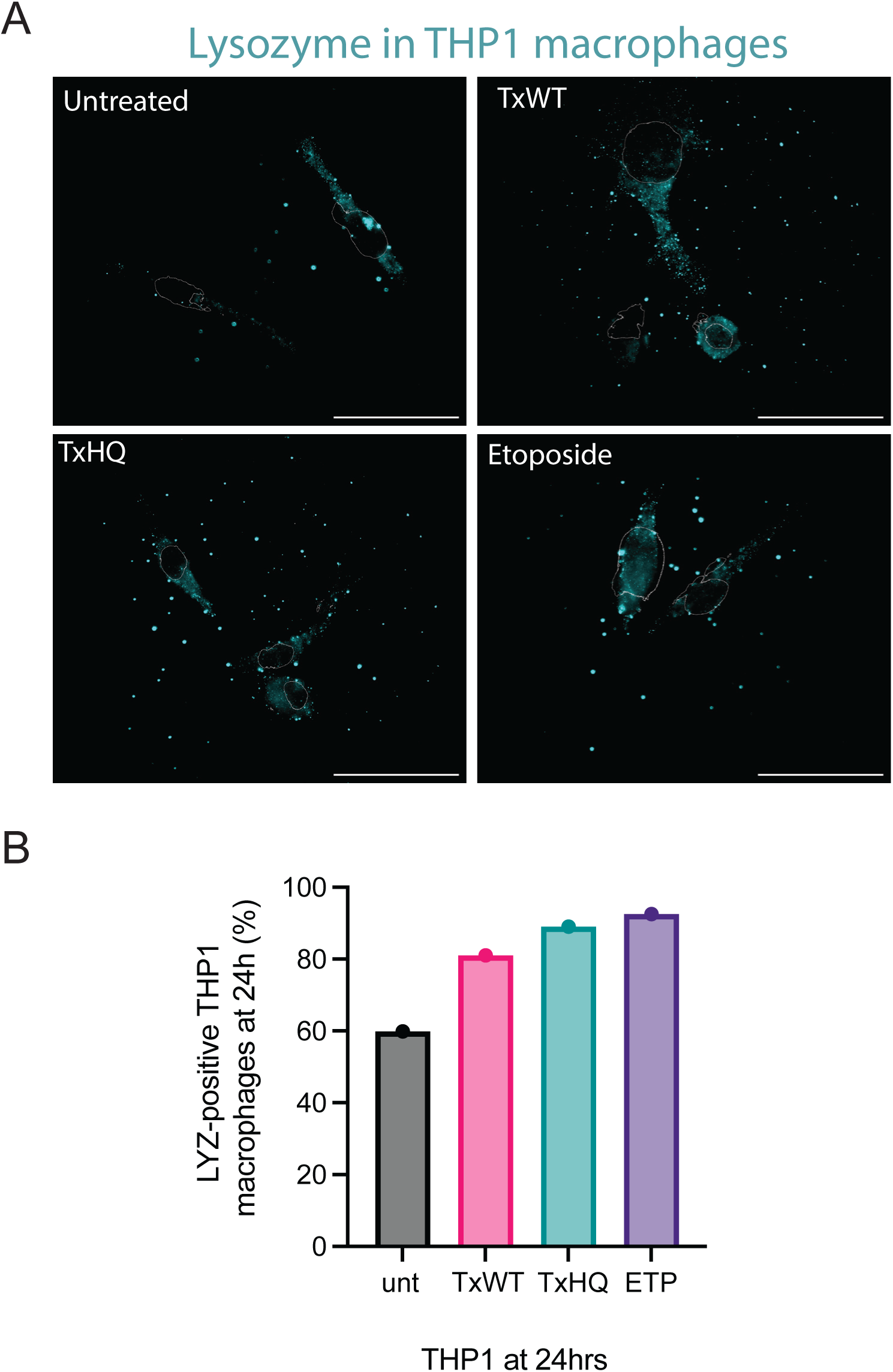
LYZ expression in THP1 macrophage cells treated with typhoid toxin. **(A)** Representative images of THP1 macrophages either untreated, treated with wild-type typhoid toxin (TxWT) or H160Q DNase-deficient toxin (TxHQ) for 2 hours prior to imaging at 24h of LYZ (cyan) or DAPI (nuclear outlines). Etoposide was used as a positive control for induction of DDRs and was incubated for the duration of the 24h experiment. Scale bars 50µm. **(B)** Bar chart showing proportion of LYZ-positive cells at 24h. Circles indicate number of biological repeats.

**Figure S4:**
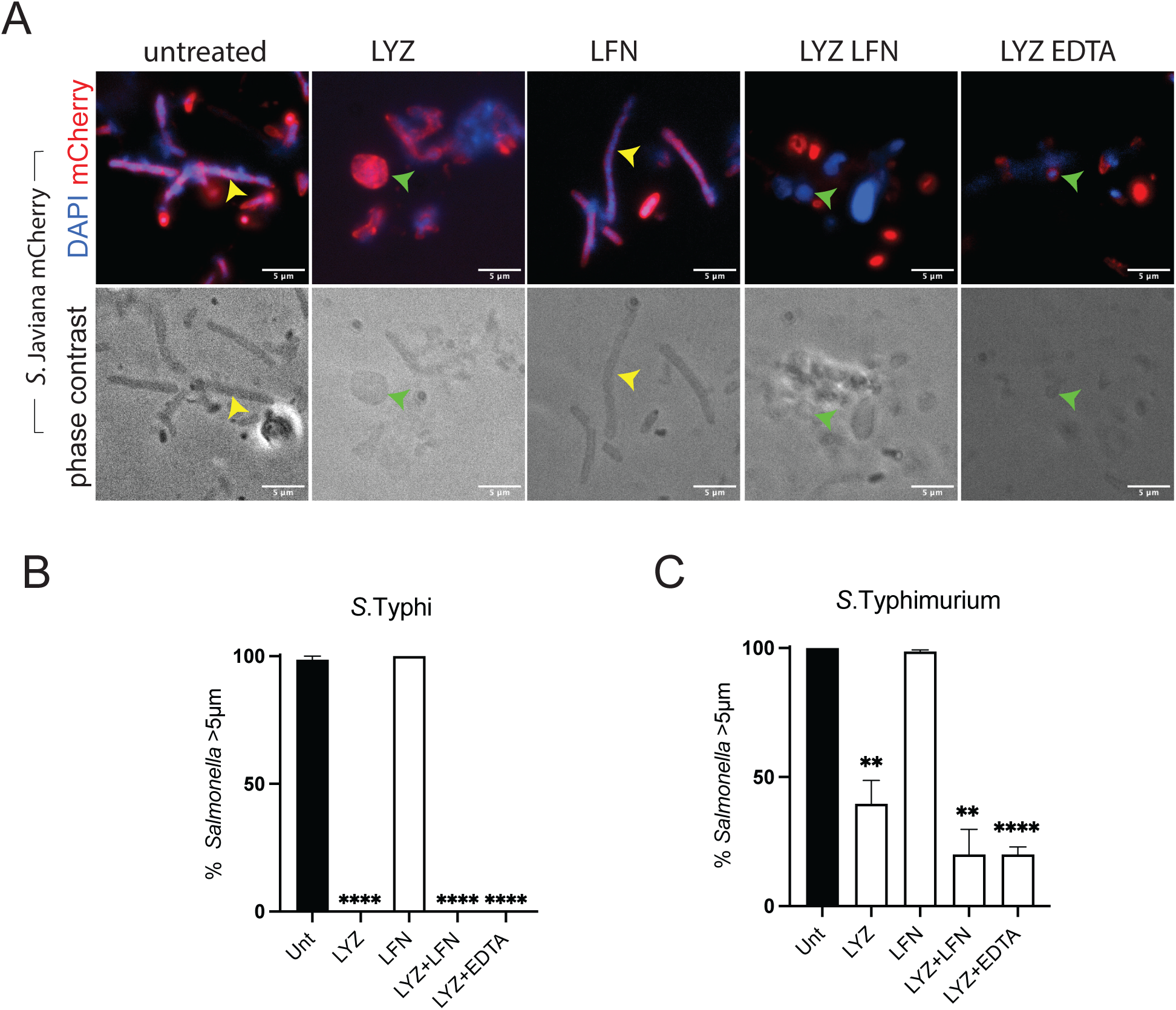
Cephalexin-treated *Salmonella* show spheroplast formation in response to LYZ. **(A)** Images of Cephalexin-treated *S*.Javiana pFPV-mCherry treated with LYZ, LFN, LYZ-LFN or LYZ-EDTA before mCherry and DAPI-stained *Salmonella* by fluorescence microscopy (top panel) or phase contrast (bottom panel). Elongated bacteria >5µm (yellow arrows) and spheroplasts (green arrows) are indicated. Scale bars 5µm. Bar charts showing the proportion of long bacteria (>5µm) following cephalexin treatment of **(B)** *S*.Typhi or **(C)** *S*.Typhimurium in the absence (unt) or presence of LYZ, LFN, LYZ and LFN, LYZ and EDTA. Asterisks in **(B), (C)** indicate significance calculated by one way ANOVA, data represented as mean ±SEM. *p<0.05, **p<0.01, ***p<0.001, ****p<0.0001.

## Notes

### Competing Interest Statement

The authors have declared no competing interest.

### Summary of Updates

To include supplementary datasets S1, S2 and their citation in the text

